# Assistive loading promotes goal-directed tuning of stretch reflex gains

**DOI:** 10.1101/2022.10.28.514224

**Authors:** Frida Torell, Sae Franklin, David W. Franklin, Michael Dimitriou

## Abstract

Voluntary movements are prepared before they are executed. Preparatory activity has been observed across the CNS and recently documented in first order neurons of the human PNS i.e., in muscle spindles. The changes seen in these sensory organs suggest that the independent modulation of stretch reflex gains may represent an important component of movement preparation. The aim of the current study was to further investigate the preparatory modulation of short- and long-latency stretch reflex responses (‘SLR’ and ‘LLR’) of the dominant upper limb. Specifically, we investigated how different target parameters (target distance and direction) affect the preparatory tuning of stretch reflex gains in the context of goal-directed reaching, and whether any such tuning depends on preparation duration and the direction of background loads. We found that target distance produced only small variations in reflex gains. In contrast, both SLR and LLR gains were strongly modulated as a function of target direction, in a manner that facilitated the upcoming voluntary movement. This goal-directed tuning of SLR and LLR gains was present or enhanced when the preparatory delay was sufficiently long (>250 ms) and the homonymous muscle was unloaded i.e., when a background load was first applied in the direction of homonymous muscle action (assistive loading). The results extend further support for a relatively slow-evolving process in reach preparation that functions to modulate reflexive muscle stiffness, likely via the independent control of fusimotor neurons. Such control can augment voluntary goal-directed movement and is triggered or enhanced when the homonymous muscle is unloaded.

**Significance Statement:** It is well-known that movement preparation improves motor performance. That is, briefly delaying the onset of a goal-directed movement can significantly benefit the overall quality of movement. However, the mechanisms underlying movement preparation remain unclear. In this study we examined the preparatory modulation of short- and long-latency stretch reflex responses in the dominant upper limb. We found that goal-directed tuning of stretch reflex gains is consistently triggered or enhanced in cases where preparation is sufficiently long (>250 ms) and a background -‘assistive’- load is first applied in the direction of homonymous muscle action. A better understanding of movement preparation will likely also benefit the development of rehabilitation regimes and movement augmentation devices.

## Introduction

Voluntary movements undergo preparation prior to their execution (Ghez et al., 1997; Kutas & Donchin, 1974; Wise, 1985). That is, voluntary movement normally involves a period of preparatory tuning where neural activity is functionally adjusted before movement initiation. This preparatory activity in the CNS correlates well with parameters such as movement direction (Tanji & Evarts, 1976), reach distance (Messier & Kalaska, 2000), movement speed (Churchland et al., 2006), reach trajectory (Hocherman & Wise, 1991) and visual target location (Batista et al., 2007). Motor preparation has long been thought of as the assembly of motor subroutines for later execution (Sternberg et al., 1978) or underlying the computation of appropriate timing and force level in the muscles tasked with achieving the motor goal (Brooks 1979). Reaching movements involve the formation of feedforward motor commands and as well as feedback policy that modulates long-latency reflex responses prior to movement onset (Ahmadi-Pajouh et al., 2012; Shadmehr & Krakauer, 2008; Todorov & Jordan, 2002; Wagner & Smith, 2008; Yeo et al., 2016).

Stretching of muscles in the upper extremities produces a ‘short latency’ reflex response (SLR) beginning approximately 20-25 ms after stretch onset, and a ‘long latency’ response (LLR) beginning 50 ms after stimulus onset (Hammond, 1956; Marsden et al., 1972). Both the SLR and the LLR are considered involuntary, with voluntary control of upper extremities beginning ~100 ms after onset of the triggering sensory event (Yang et al., 2011). It has long been known that stretch reflex responses can allow for effective resistance against unwanted postural perturbations (Nichols & Houk, 1976). However, reflex responses can also be modulated subconsciously to accommodate the execution of voluntary movements. For example, in tasks involving active reaching, significant modulation of reflex responses have been observed according to target shape (Nashed et al., 2012), obstacles (Nashed et al., 2014), static and moving targets (Cluff & Scott, 2015), and target cue direction (Pruszynski et al., 2008) (for review see (Scott, 2016)).

In the upper extremities, the SLR is often characterized by automatic gain-scaling i.e., a pre-load sensitivity, whereas the LLR is found to be robustly task-dependent (Pruszynski et al., 2009; Pruszynski et al., 2011b). More recently, the early LLR response (‘R2’) has been shown to contain a stabilizing component modulated independently of the voluntary action planned in relation to a queued target, while the late LLR (‘R3’) displayed task-dependency (Lee & Perreault, 2019). Such studies reinforce the idea that at least certain components of the LLR response are highly adaptable and not strictly “reflex” in nature (Shemmell et al., 2010). As mentioned above, goal-directed feedback controllers affecting LLR responses are thought to be already loaded during the reach preparation phase (Ahmadi-Pajouh et al., 2012). However, it is still unclear what specific mechanism produces this tuned reflex modulation prior to movement initiation.

It is currently believed that cortical preparatory activity serves a dynamic system that defines the initial conditions for the progression of goal-directed movement (Churchland et al., 2012, Churchland et al., 2010). However, exactly how cortical preparation manifests as improved motor performance is unclear. Recent findings propose one more specific neural mechanism in reach preparation. That is, movement preparation appears to include the independent and goal-directed tuning of muscle spindles, leading to a congruent modulation of reflex gains (Papaioannou & Dimitriou, 2021). These findings suggest that two independent output mechanisms are involved in the preparation of voluntary reaching i.e., one involving the direct control of α motor neurons, and another implicating independent γ motor control (Dimitriou, 2021; Dimitriou, 2022). By modulating the sensitivity of muscle spindles and stretch reflexes during preparation, the nervous system can adjust sensory feedback and reflex muscle stiffness independently of any coinciding muscle force during preparation. Our previous work has shown that target direction affects preparatory tuning of muscle spindles and stretch reflex gains in a manner that that facilitates the upcoming voluntary movement (Papaioannou & Dimitriou, 2021). In the current study, we hypothesized that both target direction and target distance may impact the preparatory tuning of stretch reflex gains. In particular, here we examine whether the fundamental parameter of target distance affects preparatory tuning of stretch reflex gains in the context of goal-directed reaching, and how any such tuning is affected by background loads, target direction and preparation duration. We found that target distance produced only small variations in reflex gains, but short and long latency reflexes were strongly tuned according to target direction, in a manner that facilitated the upcoming voluntary reach. This goal-directed tuning of stretch reflex gains was present or enhanced when the preparatory delay was sufficiently long, and the homonymous muscle was unloaded.

## Materials and Method

### Subjects

A total of 16 right-handed, neurologically healthy participants took part in the study; eight who self-identified as male (mean age 24.4 ± 4.4 years) and eight who self-identified as female (mean age 25.3 ± 6.5 years). All participants were naive as to the specific purposes of the executed task. All were financially compensated for their contribution and gave informed, written consent prior to participating in the study, per the Declaration of Helsinki. No power calculation was used to pre-determine the number of participants to include, but we used a similar or larger number of participants than previous studies (e.g., Dimitriou et al., 2012; Pruszynski et al., 2009; Weiler et al., 2021; Yang et al., 2011). Two of the 16 participants did not conform to the experimental manipulations of the study (i.e., presented excessive contraction in shoulder muscles in conditions requiring muscle relaxation/unloading), and their data were therefore not included in further analyses. The current experiments were part of a research program approved by the Ethics Committee of Umeå University, Umeå, Sweden.

### The experimental setup

Participants sat upright in a customized adjustable chair in front of the Kinarm robotic platform (Kinarm end-point robot, BKIN Technologies Ltd., Canada). The participants used their right hand to grasp the robotic manipulandum (Fig. 1A). The right forearm was placed inside a customized foam structure, resting on an airsled that allowed frictionless movement of the arm in a 2D plane. To ensure a secure mechanical connection between the forearm, foam-cushioned airsled, the Kinarm^TM^ handle and the hand were secured using a leather fabric with Velcro attachments. This attachment also fixated the wrist in straight alignment with the forearm throughout the experiment. The robotic platform measured kinematic data regarding the position of the hand, and sensors inside the robotic handle recorded the forces exerted by the participants’ right hand (six-axis force transducer; Mini40-R, ATI Industrial Automation, U.S.A.). Position and force data was sampled at 1 kHz.

**Fig. 1.**
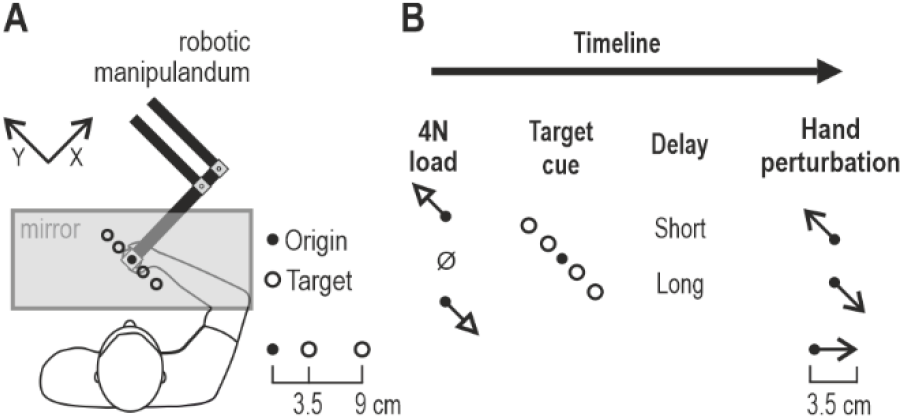
The robotic platform and experimental setup. **(A)** Participants manipulated the position of a robotic handle using their right hand. The right forearm rested on an airsled that allowed frictionless movement. All visual stimuli were projected onto a one-way mirror. The participant could not view their hand or the robotic handle, and the position of the hand was represented by a visual cursor in the plane of movement. The potential visual targets were placed along the Y-axis (defined as illustrated). Specifically, two ‘near’ and two ‘far’ targets were used, placed at 3.5 cm and 9 cm from the origin, respectively. **(B)** The task timeline. Each trial was initiated when the participant kept the hand/cursor immobile at origin. Either no load was applied or a slow-rising 4N load in either the +Y (upper left direction) or -Y direction (lower right direction) was then applied. Regardless of load, the participants had to keep their hand at origin. A target was then cued by turning red, and remained in this state for either a relatively short (250 ms) or long (750 ms) delay period. This preparatory delay was followed by a rapid −150 ms- perturbation of the hand by 3.5 cm in either the +Y or -Y direction. The cursor position was frozen during the 150 ms perturbation. At the end of the perturbation, the red target turned green (‘Go’ signal), and the participants were instructed to rapidly complete movement to the target. All trials were block-randomized, meaning that the direction of the kinematic perturbation was unpredictable, even after experiencing a specific load, target cue and delay.

### Experimental design

Participants viewed the content of a monitor through the reflection of a one-way mirror which prevented view of the hand. The position of the right hand was represented by a white dot (‘cursor’; 1 cm diameter) in the plane of movement. The origin was represented by a circle of 1.3 cm diameter.Four visual targets in total were placed along the Y axis (see Fig. 1A). Specifically, two targets were placed in front-and-left direction (‘+Y’) and two in right-and-back direction (‘−Y’). The size of all targets was the same (2.4 cm diameter). The center-to-center distance between the origin and each ‘near’ target was 3.5 cm, and 9 cm between the origin and each ‘far’ target (Fig. 1A). All potential targets were continuously displayed as orange circle outlines.

The participant begun each trial by moving the cursor/hand inside the origin circle. There, they had to remain immobile for a random wait period ranging from 1-1.5 seconds. At this point either no load was applied, or controlled forces could then be applied in the form of a slow-rising 4 N load in either the +Y or −Y direction (800 ms rise-time and a 1200 ms hold). After the hold phase, one of the four targets then became a red filled circle, representing the target cue. The participant had to remain at the origin for a preparatory delay period of 250 ms or 750 ms, also referred to as the relatively ‘short’ and ‘long’ delay, respectively. The two preparatory delays were chosen based on previous results, using a similar setup (Dimitriou, 2018; Papaioannou & Dimitriou, 2021). At the end of the preparatory period, a position-controlled perturbation displaced the hand by 3.5 cm in either the +Y or −Y direction (rise time of 150 ms, no hold). The cursor position was frozen at origin during the 150 ms perturbation. The haptic perturbations were used to produce displacements with approximate bell-shaped velocity profiles. The KINARM robot was able to exert the appropriate stiffness (maximum ~40,000 N/m), regardless of load/force conditions, to ensure the desired hand kinematics on every trial. Upon the end of the perturbation, the cued target (red filled circle) turned green (representing the ‘Go’ signal), and the participants were instructed to complete the movement to the target. In other words, the ‘Go’ signal to reach the target was always given upon the end of the brief position perturbation.

Once the cursor/hand reached the target and remained there for 300 ms, the trial ended, and participants were given visual feedback on their performance. The performance metric measured the time from the ‘Go’ signal until the target was reached. A time faster than 400 ms resulted in a ‘Too fast’ message being shown on the monitor, 400-1400 ms resulted in ‘Correct’ and >1400 ms in ‘Too Slow’. The chosen feedback intervals motivated rapid goal-directed behaviour (see e.g., Figs 2–3) but also allowed sufficient time for participants to reach the goal regardless if the hand was first perturbed in an incongruent direction. After receiving visual feedback (lasting for 300 ms), the participants returned the cursor inside the origin in order to start the next trial. After any trial, the participant could move the cursor to the side of the workspace and rest, although breaks were normally encouraged and requested after a substantial number of trials. Breaks normally lasted for <5 minutes. The experiment consisted of 48 unique trial types: 4 targets (+Y far, +Y near, −Y far, and −Y near) x 3 load conditions (4N in +Y direction, null-load, and 4N in −Y direction) x 2 delay durations (250 ms and 750 ms) x 2 perturbation directions (+Y and −Y). The participants performed 15 repetitions of each condition i.e., the total number of trials in each experiment was 720. The trials were presented in a block-randomized order, where one ‘block’ represented a set of 48 unique trials. The delayed-reach task took approximately 1.5 hours to complete.

**Fig. 2.**
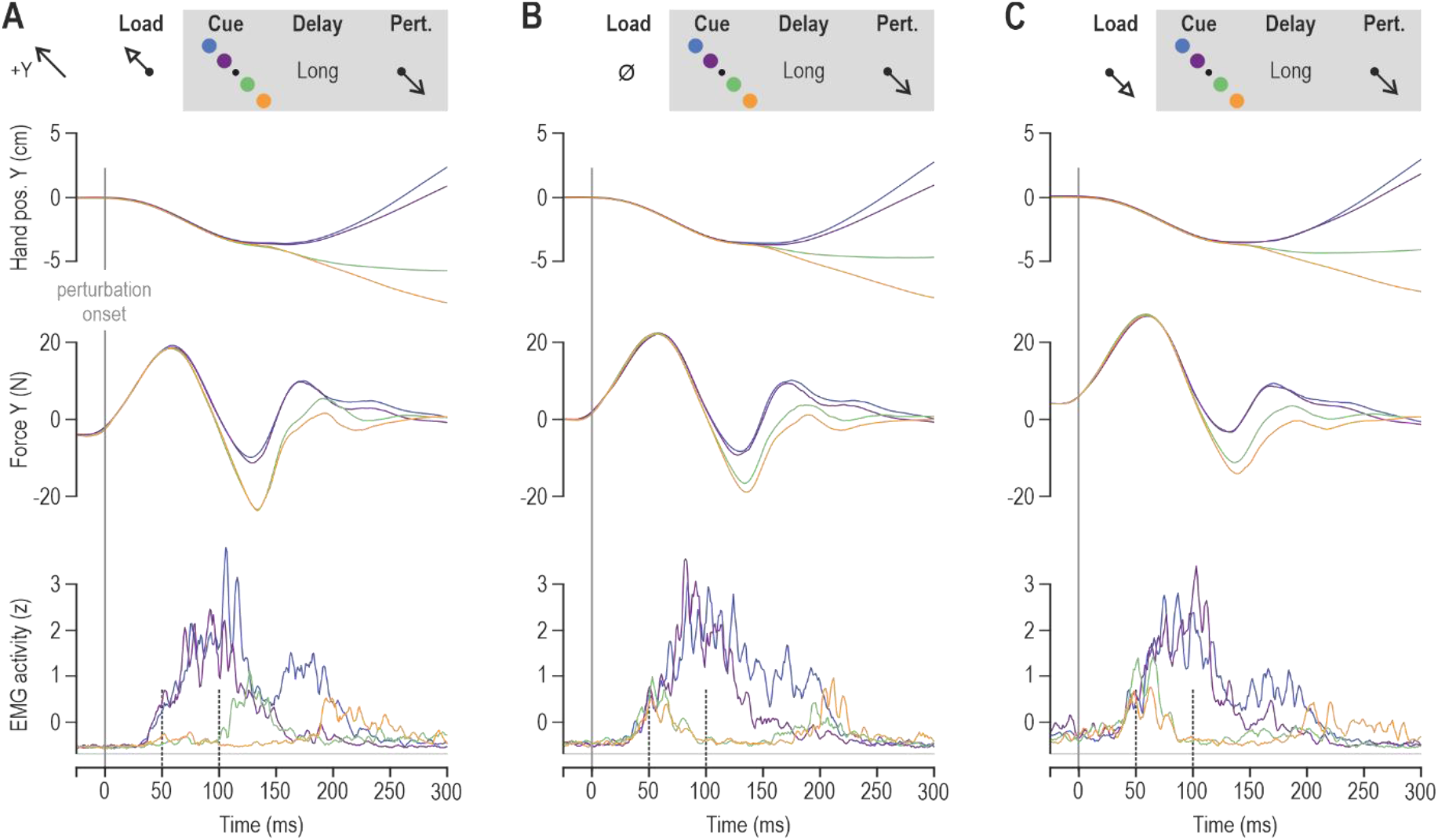
Goal-directed responses from the pectoralis of a single participant in the delayed reaching task. Throughout, blue and purple traces represent trials where participants had to reach for the ‘far’ and ‘near’ targets in the +Y direction, respectively (see also schematics). Reaching these targets required shortening of the pectoralis. Green and orange traces represent trials where the participant had to reach the ‘far’ and ‘near’ targets in the −Y direction, respectively. These trials require stretch of the pectoralis. All data in this figure involved a long preparatory delay (i.e., 750 ms) and are aligned on perturbation onset (time zero). **(A)** A slow-rising load (‘pre-load’) was first applied in the +Y direction before the visual target cue and subsequent haptic perturbation that stretched the pectoralis. **(B)** As ‘A’ but no slow-rising load was applied. **(C)** As ‘A’ but the slow-rising load was applied in the −Y direction, loading the pectoralis before the visual target cue and subsequent haptic perturbation.

**Fig. 3.**
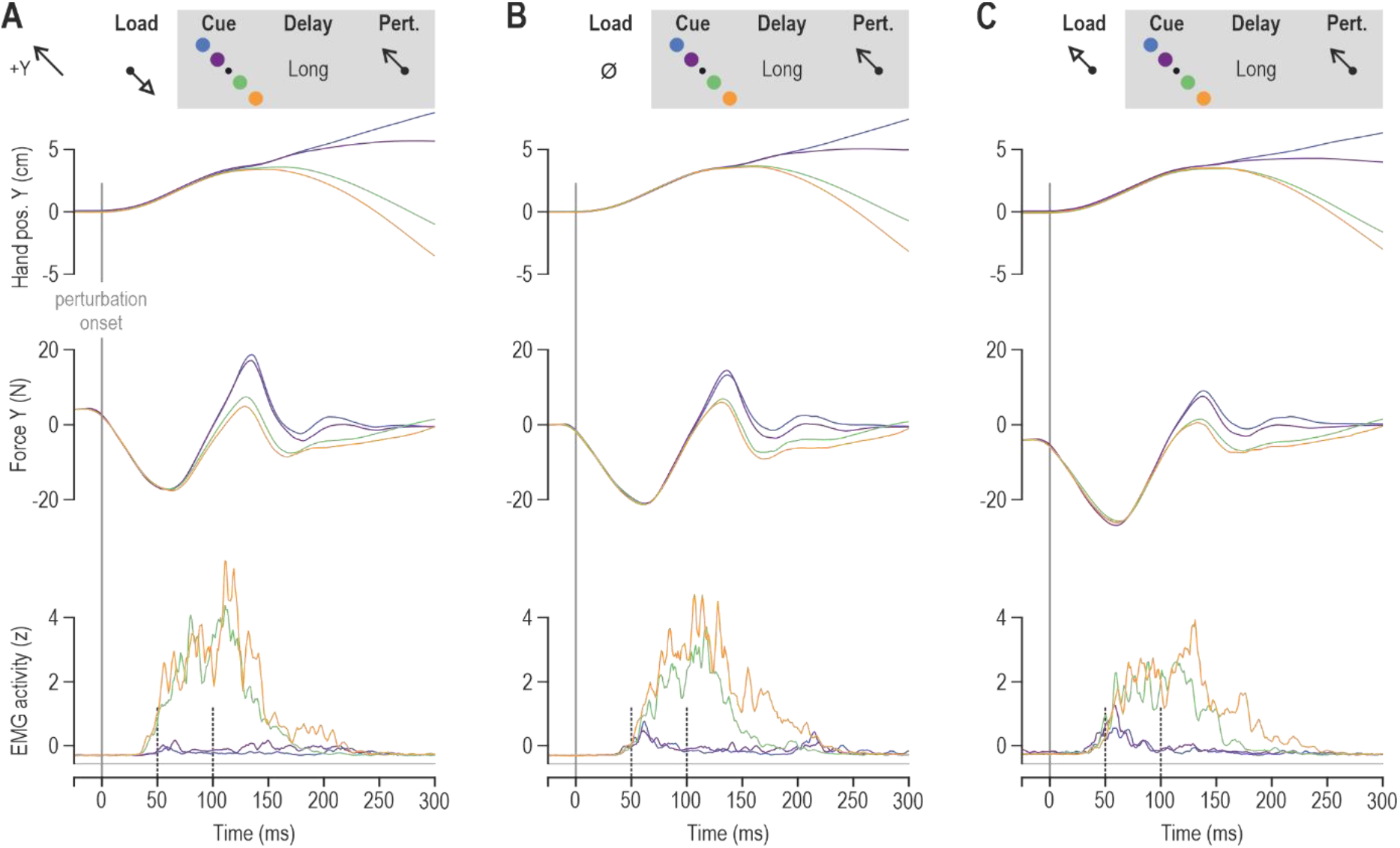
Goal-directed responses from the posterior deltoid of a single participant in the delayed reaching task. Throughout, blue and purple traces represent trials where participants had to reach for the ‘far’ and ‘near’ targets in the +Y direction, respectively. Reaching these targets required stretch of the posterior deltoid. Green and orange traces represent trials where the participant had to reach the ‘far’ and ‘near’ targets in the −Y direction, respectively. These trials required shortening of the posterior deltoid. All data in this figure involved a long preparatory delay (i.e., 750 ms) and are aligned on perturbation onset (time zero). **(A)** A slow-rising load (‘pre-load’) was first applied in the −Y direction before the visual target cue and subsequent haptic perturbation that stretched the posterior deltoid. **(B)** As ‘A’ but no slow-rising load was applied. **(C)** As ‘A’ but the slow-rising load was applied in the +Y direction, loading the posterior deltoid before the visual target cue and subsequent haptic perturbation.

### Electromyography (EMG)

Surface electromyography (EMG) signals were recorded from (1) m. brachioradialis, (2) m. biceps brachii, (3) m. triceps brachii caput laterale, (4) m. triceps brachii caput longum, (5) m. deltoideus pars anterior (6) m. deltoideus pars posterior, and (7) m. pectoralis major. However, in line with previous findings, it was the latter three muscles (i.e., shoulder actuators: pectoralis, anterior and posterior deltoid) that were primarily engaged in the delayed reach task of our setup, hence only these three muscles were used for statistical analyses. We used surface electromyography (EMG) electrodes (Bagnoli^™^ DE-2.1, Delsys Inc., USA) that have contact dimensions 10.0 x 1.0 mm with 10 mm inter-electrode spacing. Prior to attaching the electrodes, the skin was cleaned using alcohol swabs. The electrodes were coated with conductive gel and placed on the peak of the belly of the studied muscles in the direction of the muscle fibres. All electrodes were attached with double-sided tape and further secured using surgical tape. One ground electrode (Dermatrode^®^ HE-R Reference Electrode type 00200-3400; American Imex, Irvine, CA, USA), with a diameter of 5.08 cm, was placed on the processus spinosus of the C7 region. The EMG signals were analogue band-pass filtered through the EMG system (20 – 450 Hz) and sampled at 1 kHz.

### Data preprocessing

EMG data was high pass filtered using a fifth-order, zero phase-lag Butterworth filter with a 30 Hz cut-off and then rectified. For each trial, onset of movement was defined as the point where velocity first reached 5% of peak velocity during movement. Normalization was applied to allow EMG data from different muscles and participants to be combined and/or compared. The raw data was normalized (z-transformed) using a procedure that has been described in more detail elsewhere (Dimitriou, 2014; Dimitriou, 2016). Briefly, this involves concatenating all EMG data (here, from all 15 blocks) and calculating a grand mean and grand standard deviation for each muscle separately. Normalized EMG data for each muscle is obtained by subtracting the grand mean and dividing it by the grand standard deviation. The first five blocks of trials were viewed as familiarization trials and were not included in the analyses. The focus of this study was on stretch reflex responses, therefore we only analysed data from stretching muscles. That is, we analysed specific combinations of muscle and perturbation direction. To simplify analyses of individual muscles, the median EMG signal of each muscle was generated for each trial type (i.e., averaged for each load, perturbation, preparatory delay, target direction and distance) for each participant. Data pre-processing was performed using MATLAB^®^ (MathWorks, 2020b, Natick, MA, USA). For plotting purposes only, the EMG signals were smoothed using a 5 ms moving window.

### Statistics

Statistical analyses were done on the z-normalized EMG data, available at Mendeley Data, V1, doi: 10.17632/hnfp5yrght.1. To check normality Shapiro-Wilks test for samples with <50 data-points and Lilliefors test for larger samples were used. For each muscle, the data used for statistical analyses were the averages across the pre-perturbation epoch (i.e., period starting 20 ms prior to perturbation onset), the SLR epoch (25-50 ms post-perturbation onset) and the ‘late’ LLR epoch (76-100 ms). To analyse pre-perturbation and reflex EMG responses, a repeated-measures analysis of variance (ANOVA) of the design 2 (preparatory delay) x 3 (load) x 2 (target direction) × 2 (target distance) was used. For the SLR response in particular, the pre-loaded (‘loaded’) muscle condition was analysed separately (i.e., 2 delay x 2 distance x 2 direction), as it is well known that automatic gain-scaling due to muscle pre-loading tends to saturate the SLR response, preventing its goal-directed modulation, as occurred in our study (e.g., see Figures 2–6 and Results). Throughout, post-hoc analyses were performed using Tukey’s HSD (honest significant difference).

**Fig. 4.**
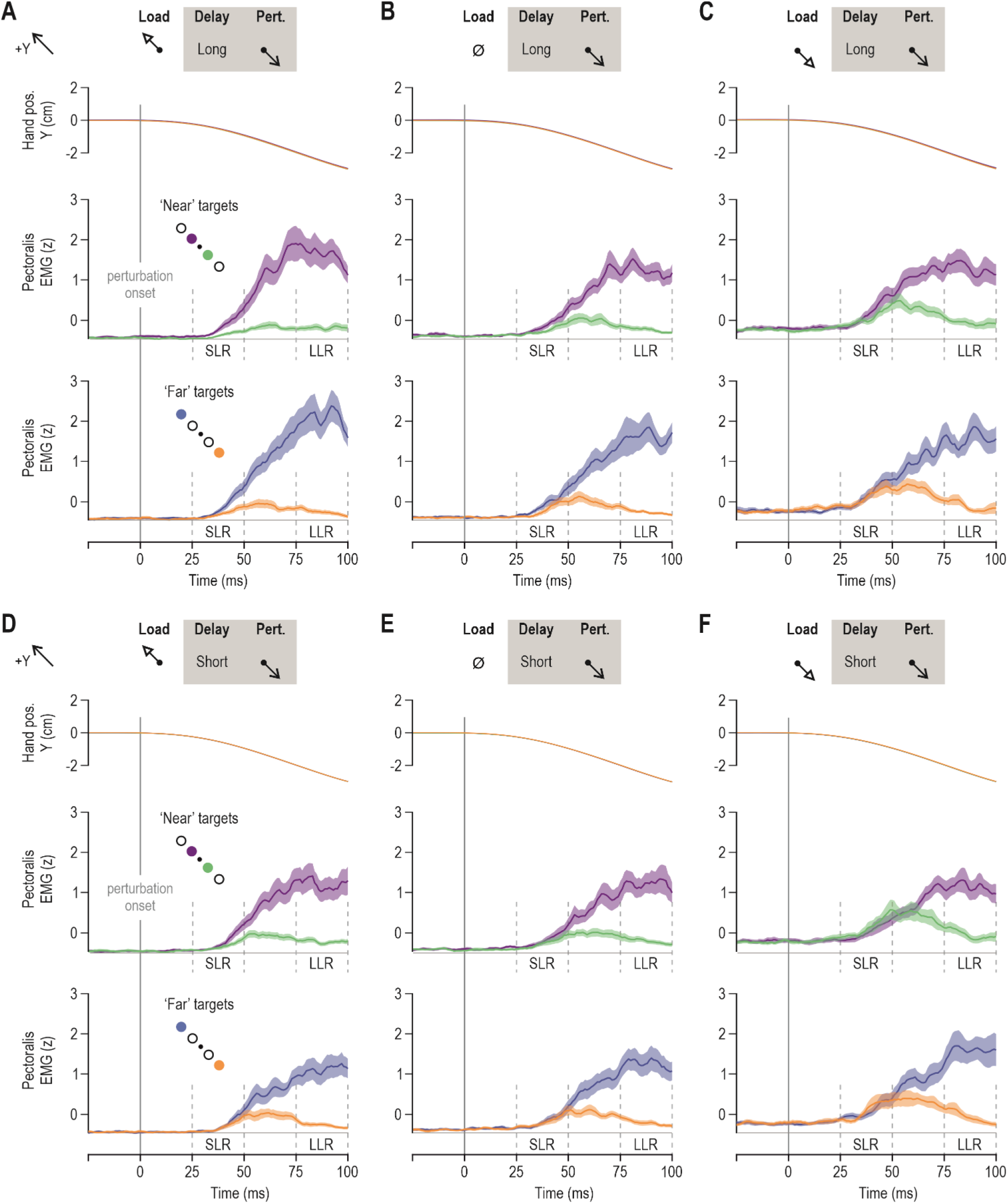
Goal-, load- and delay-dependent stretch reflex responses of the pectoralis. Colour-coding as in previous figures. **(A)** The upper panel represents mean hand position across participants (N = 14), for all trials where the pectoralis muscle was unloaded before being stretched by the perturbation, following a long preparatory delay (750 ms). The middle row displays mean pectoralis EMG activity across participants for the subset of trials where one or the other ‘near’ targets were cued for a long delay before pectoralis stretch; the bottom panel represents the equivalent for ‘far’ targets. **(B)** As ‘A’, but representing the ‘no load’ trials **(C)** As ‘A’ but representing trials where the pectoralis was loaded before the stretch perturbation. **(D-F)** As ‘A-C’ but representing trials where the preparatory delay was short. See also schematics. Throughout, colour shading represents ±1 SEM.

**Fig. 5.**
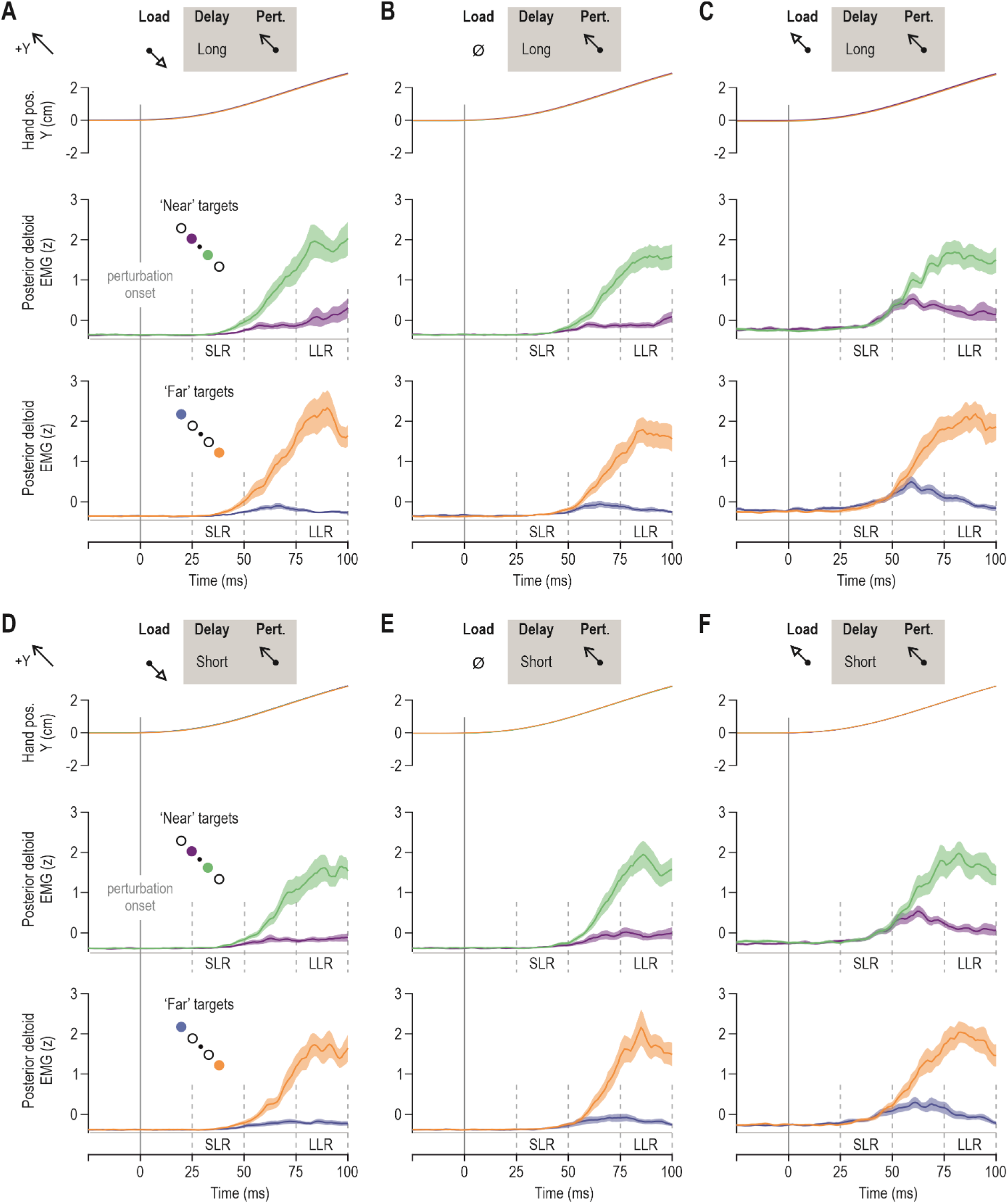
Goal-, load- and delay-dependent stretch reflex responses of the posterior deltoid. Colour-coding as in previous figures. **(A)** The upper panel represents mean hand position across participants (N = 14), for all trials where the posterior deltoid was unloaded before being stretched by the perturbation, following a long preparatory delay (750 ms). The middle row displays mean posterior deltoid EMG activity across participants for the subset of trials where one or the other ‘near’ targets were cued for a long delay before posterior deltoid stretch; the bottom panel represents the equivalent for ‘far’ targets. **(B)** As ‘A’, but representing the ‘no load’ trials. **(C)** As ‘A’ but representing trials where the posterior deltoid was loaded before the stretch perturbation. **(D-F)** As ‘A-C’ but representing trials where the preparatory delay was short. See also schematics. Throughout, colour shading represents ±1 SEM.

**Fig. 6.**
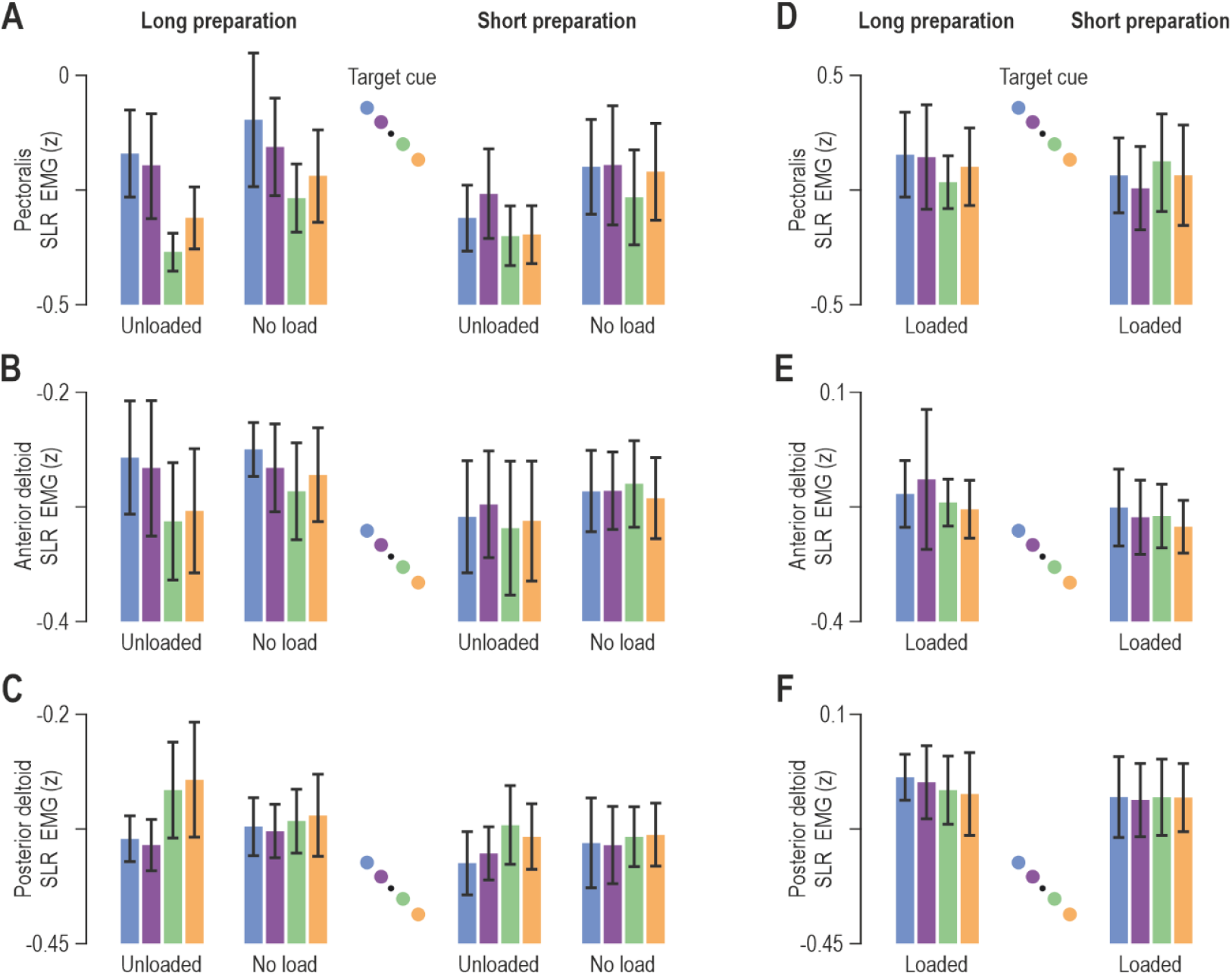
Goal-directed modulation of short-latency reflex gains. **(A)** The coloured bars represent mean pectoralis SLR EMG (z) across participants (N=14), and vertical lines represent 95% confidence intervals. Colour-coding as in previous figures (see also schematic). Data originate from trials where the homonymous muscle was unloaded and when there no pre-load, as indicated. An ANOVA indicated a consistent effect of target direction on pectoralis SLR gains when the preparatory delay was long (left columns). That is, SLR gains are relatively suppressed when allowed long-enough time to prepare reaching a target along the direction of homonymous muscle stretch (green/orange). **(B)** As ‘A’ but representing the anterior deltoid muscle. Similar SLR modulation patterns were observed as for the pectoralis (see Results section for more details). **(C)** As ‘A’ but representing the posterior deltoid. **(D-F)** As A-C but representing trials where the homonymous muscle was loaded.

To estimate the onset of SLR reflex modulation, we used the receiver-operator characteristic (ROC) technique (Green & Swets, 1966). A ROC area of 1 and 0 signifies perfect discrimination while a ROC area of 0.5 signifies a discrimination performance equal to chance. For this type of analysis, we only used data from trials where the preparatory delay was long. Specifically, the EMG curves of targets in the direction of homonymous muscle stretch were contrasted to EMG responses observed in trials where the cued target was in the direction of muscle shortening. The obtained averages across target distance were together viewed as representing the reflex modulation in the population sample. Discrimination was viewed as significant when the ROC area remained >0.75 for five consecutive samples (Corneil et al., 2004). To assess the reflex modulation onset for each participant separately, the same type of procedure was performed using individual EMG responses across trials. All SLR modulation onsets were confirmed by visual inspection, to eliminate the risk of false positives.Data tabulation was performed using MATLAB^®^ (MathWorks R2020b, Natick, MA, USA), and statistical analyses using STATISTICA^®^ (StatSoft Inc, USA).

## Results

Participants were asked to hold their right hand within a starting position while one of three different loads was applied (−4N, 0N or 4N). One of four targets was then cued (turned red), and after either a short or long delay the limb was perturbed by 3.5 cm in one of two directions (+Y or −Y; Fig. 1A). After the 150 ms perturbation, the target colour was changed from red to green, indicating that the participants should move the hand to this target. Here we examine how the target location (direction and distance), background load and delay between the presentation of the target and the perturbation effected the reflex responses prior to the movement of the participants.

Representative data from single participants are presented in Figure 2 and 3, for the pectoralis major and posterior deltoid, respectively. Visual inspection of both figures points to differences in EMG during the SLR epoch as a function of target cue, particularly when the muscles were unloaded (Fig 2A, Fig 3A). Specifically, the SLR response appears to be relatively suppressed when the cued target is in the direction of muscle stretching (e.g., blue/purple vs green/yellow for the pectoralis; Fig 2A). Relative suppression of stretch reflexes -and therefore of muscle stiffness- would facilitate the subsequent reaching movement. Clear goal-directed differences in the LLR epoch can be seen across all load conditions (Figs 2–3). As elaborated in the Methods section, the modulation of stretch reflex responses was analysed using averaged data across participants. Figure 4 shows the averaged hand position and pectoralis EMG activity aligned on perturbation onset (time zero), whereas Figure 5 displays equivalent responses from the posterior deltoid.

### The pre-perturbation epoch

To confirm the lack of goal-directed differences in the pre-perturbation epoch (i.e., 20 ms period before perturbation onset), an ANOVA of the design 2(preparatory delay) x 3(load) x 2(target distance) x 2(target direction) was performed using averaged EMG data over this epoch. As expected, for all investigated muscles, there was a significant main effect of load condition on pre-perturbation EMG. Specifically, for the pectoralis, ANOVA indicated a significant main effect of load (F_2,26_=41.5, p<10^-5^ and 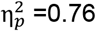), whereas all other main and interaction effects were not significant (all p>0.065). Tukey’s HSD test showed that pre-perturbation EMG was significantly higher only in the ‘loaded’ condition (vs. ‘no load’ and ‘unloaded’ conditions: all p<0.0002; Fig. 4C & F). As mentioned above, the plots point to a particularly clear effect of target direction on reflex responses when the homonymous muscle is unloaded (e.g., Fig. 4A); any equivalent differences in pre-perturbation EMG in the ‘unloaded’ condition could possibly account for the enhanced SLR effects due to gain-scaling. However, a planned comparison test indicated no impact of target direction on pre-perturbation EMG when the pectoralis was unloaded (p=0.63). The same results were obtained for the anterior deltoid. That is, the ANOVA produced a significant main effect of load (F_2,26_=5.3, p=0.012 and 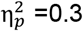) whereas all other main and interaction effects were not significant (all p>0.11). Tukey’s HSD test indicated significantly higher pre-perturbation EMG only when the muscle was loaded (vs. unloaded: p=0.016; vs. no load: p=0.035). A planned comparison also indicated no specific effect of target direction on pre-perturbation EMG when the anterior deltoid was unloaded (p=0.36).

For the posterior deltoid, an ANOVA again showed a main effect of load (F_2,26_=57.8, p<10^-5^ and 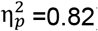) as with the other two muscles (i.e., higher EMG in ‘loaded’ condition, all p<0.0002). But there was also a main effect of preparatory delay, with generally higher pre-perturbation EMG observed when the preparatory delay was long (F_1,13_=18.4, p=0.0009 and 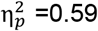). ANOVA also indicated significant interaction effects, such as between delay, load condition and target direction (F_2,26_=6.1, p=0.007 and 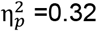). However, as can be appreciated by visually inspecting the pre-perturbation epochs in Figure 5, differences in posterior deltoid EMG as a function of target parameters were apparent only when the muscle was loaded. Indeed, performing the ANOVA analyses without including the ‘loaded’ condition eliminated all effects involving target parameters (all p>0.05); only a significant main effect of preparatory delay remained (F_1,13_=14.8, p=0.002 and 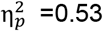), indicating higher pre-perturbation activity in the posterior deltoid following a long delay, irrespective of target direction or distance. Furthermore, a planned comparison test indicated no significant effect of target direction on pre-perturbation EMG when the posterior deltoid was unloaded (p=0.59). Importantly, there are no consistent target-dependent differences in the pre-perturbation epoch for the posterior deltoid, anterior deltoid, and pectoralis muscles of the dominant upper limb.

### The short-latency reflex epoch (SLR)

As the main aim of this study was to further investigate the preparatory modulation of stretch reflex gains, we focus the rest of the analysis on the specific responses during muscle stretch. Specifically, all data originates from trials where the hand was perturbed along the direction of targets associated with stretch of the homonymous muscle, regardless of voluntary movement intent. Moreover, it is well known that automatic gain-scaling due to pre-loading of the muscle tends to saturate the SLR response, limiting its goal-directed modulation (see Figures 2–5). Therefore, SLR responses from the pre-loaded (‘loaded’) homonymous muscles were analysed separately, using an ANOVA design of 2(preparatory delay) x 2(target distance) x 2(target direction). Analysis of SLR responses associated with the ‘no load’ and ‘unloaded’ conditions was performed using an ANOVA design of 2(preparatory delay) x 2(load) x 2(target distance) x 2(target direction). As the main aim of this study investigate the goal-directed modulation of stretch reflexes, the following text focuses on and first describes the results involving the ‘unloaded’ and ‘no load’ conditions. Throughout, differences in the stretch reflex responses – at all latencies – represent differences in stretch reflex gains, as both the initial position and kinematic perturbation of the hand are matched across relevant experimental conditions.

For the pectoralis SLR (Fig. 6A), ANOVA indicated a significant main effect of target direction (F_1,13_=12.1, p=0.004 and 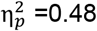), with stronger responses observed when the target cue was placed along the direction of pectoralis shortening i.e., the +Y direction. There was also a main effect of preparatory delay (F_1,13_=5.3, p=0.04 and 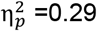) and an interaction effect between preparatory delay and target direction (F_1,13_=8.5, p=0.012 and 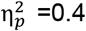). Tukey’s HSD test indicated that the impact of target direction (blue/purple > orange/green, Fig 6A) was significant only following a long preparatory delay (all p<0.008: mean blue/purple −0.18 ± 0.09 95%CI vs. mean orange/green −0.29 ± 0.09 95%CI). However, a planned comparison analysis revealed an additional significant effect of target direction on the SLR responses of the unloaded muscle following a short preparatory delay (i.e., purple > green, p=0.036: mean purple −0.27 ± 0.09 95%CI vs. mean green −0.35 ± 0.08 95%CI); but, this effect was less pronounced than the equivalent following a long delay (Fig. 4A vs. Fig. 4D). There was also a main effect of target distance on pectoralis SLR, with stronger overall responses evident for ‘far’ vs. ‘near’ targets (F_1,13_=9.2, p=0.0094 and 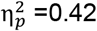). The interaction effect between preparatory delay and target distance failed to reach significance (F_1,13_=4.4, p=0.056 and 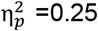) Interestingly, there was also a main effect of load condition on pectoralis SLR, with stronger EMG responses in the ‘no load’ vs. ‘unloaded’ condition (F_1,13_=8, p=0.014 and 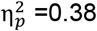).

Similar results were obtained with regard to the SLR responses of the anterior deltoid (Fig. 6B). Specifically, an ANOVA indicated a significant main effect of target direction (F_1,13_=11.6, p=0.005 and 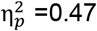), with stronger SLR responses when the target cue was placed along the direction of muscle shortening. The ANOVA also revealed a main effect of preparatory delay on anterior deltoid SLR, with the long delay associated with higher responses (F_1,13_=8.6, p=0.012 and 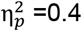). For the same muscle, there was also a significant interaction effect between preparatory delay and target direction (F_1,13_=5.2, p=0.04 and 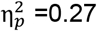). Tukey’s HSD test indicated significantly higher SLR responses in the long delay condition when the target cue was in the direction associated with shortening of the anterior deltoid (all p<0.008). However, there was no main effect or interaction effect involving target distance for this muscle (all p>0.5). Overall, for both the anterior deltoid and pectoralis, there was a clear goal-directed modulation of SLR gains in cases where a long-enough time (>250 ms) was allowed for preparing stretch of the homonymous muscle (Fig 6A&B, far left columns).

With regard to the SLR responses of the posterior deltoid (Fig. 6C), an ANOVA indicated both a main effect of preparatory delay (F_1,13_=12.7, p=0.0034 and 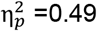) and target direction (F_1,13_=7.2, p=0.019 and 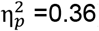), with higher SLR EMG for longer preparatory delays and for target cues presented along the direction of muscle shortening i.e., −Y direction for the posterior deltoid. There was also an interaction effect between load and target direction (F_1,13_=5.4, p=0.037 and 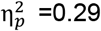), with post-hoc analyses indicating a significant impact of target direction on posterior deltoid SLR only in the ‘unloaded’ condition (p=0.0048: mean orange/green −0.28 ± 0.05 95%CI vs. mean blue/purple −0.34 ± 0.02 95%CI). For this muscle, there was also a significant interaction effect between preparatory delay and target distance (F_1,13_=8.4, p=0.012 and 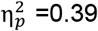) although Tukey’s HSD test indicated no differential impact of target distance as a function of preparatory delay (Fisher’s test on the other hand indicated a significant impact of target distance when the delay was long; p=0.026). Overall, we again find a clear target-related modulation of the SLR in the unloaded condition when sufficient preparation time is provided (Fig 6C, far left column).

Figure 6D–F displays SLR responses across participants when the homonymous muscle was loaded. ANOVA analyses (2 delay x 2 target distance x 2 target direction) indicated no significant main effect (or interaction effect) of target location on the SLR responses of the three analysed muscles (all p>0.28 for anterior deltoid; all p>0.1 for pectoralis; all p>0.39 for posterior deltoid). For the loaded anterior deltoid alone (Fig. 6E), there was a significant main effect of preparatory delay on SLR responses (long>short), with (F_1,13_=7.95, p=0.014 and 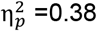), whereas no such effect was observed for the posterior deltoid and pectoralis (p=0.15 and p=0.14, respectively). The above analyses confirm what can be visually appreciated by inspecting SLR responses in Figures 2–6: loading the homonymous muscle substantially for the purposes of postural maintenance tends to saturate the SLR response, thwarting its preparatory modulation according to task goals.

In summary, across the three analysed muscles, ANOVA tests yielded no consistent effect of target distance on SLR responses. That is, an effect of target distance on SLR (‘far’ > ‘near’ targets) was evident only for the pectoralis, if the muscle was not first loaded. In contrast, there was a consistent effect of target direction. Specifically, all muscles produced goal-directed SLR responses, in the sense of displaying relatively weaker/stronger reflex gains, when preparing to reach targets requiring stretch/shortening of the homonymous muscle. These response patterns were present or clearer when the preparation time was sufficiently long and the muscles were unloaded i.e., when a background load was first applied in the direction of muscle action (Fig 6, far left columns).

To further characterise the goal-directed modulation of the SLR, receiver operator characteristic (ROC) analysis was applied. Since ANOVA indicated no consistent effect of target distance on SLR responses, the data was collapsed across target distance, to concentrate on the impact of target direction. The relevant difference signals were created by contrasting the EMG curve observed when reaching for targets requiring muscle stretch vs. when reaching for targets requiring muscle shortening were examined to determine the time point at which the signals could be discriminated by an ideal observer. For the unloaded pectoralis (Fig. 7A), dog leg fits indicated deviance at 19 ms, for the no load condition at 23 ms and for the loaded condition at 34 ms. For the unloaded posterior deltoid (Fig. 7B), this occurred at 28 ms, for the no load condition at 41 ms and for the loaded condition at 50 ms. The onset times were also calculated for each participant individually (small red circles in Fig. 7A–B). The time point at which the ROC was above 0.75 was also identified. For the unloaded pectoralis (Fig. 7A), this occurred at 45 ms, for the no load condition at 55 ms and for the loaded condition at 57 ms. For the unloaded posterior deltoid (Fig. 7B), this occurred at 52 ms, for the no load condition at 61 ms and for the loaded condition at 63 ms (red vertical lines).

**Fig. 7.**
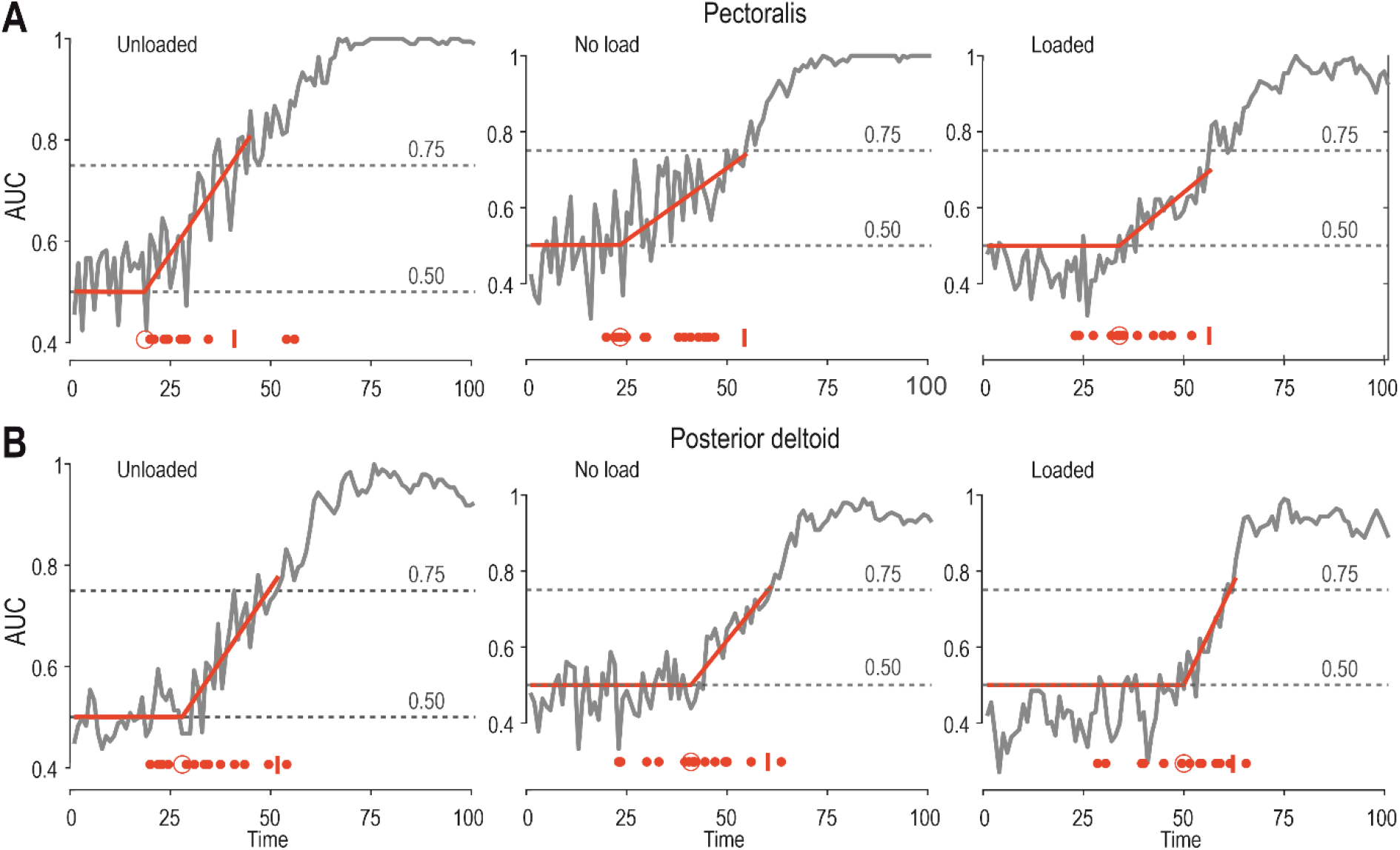
The time onset of SLR modulation. **(A)** The grey curve in each panel represents the area under the ROC, pertaining to pectoralis SLR modulation as a function of target direction, after experiencing one of the three load conditions (‘unloaded’, ‘no load’ and ‘loaded’) and a long preparatory delay (see Methods and Results for more details). Specifically, vertical axes represent the probability that an ideal observer could discriminate between the EMG difference curves. Each solid red line represents a dog leg fit which was applied to determine the onset of significant SLR modulation as a function of target direction (see also larger red circle at the bottom of each panel). The small red vertical line at the bottom of each panel represents the time point when the ROC area remained >0.75 for five consecutive time points (i.e., five consecutive ms). The smaller red dots represent the ROC result for each individual participant. **(B)** As ‘A’ but representing the posterior deltoid.

### The long-latency reflex epoch ‘LLR’

The modulation of LLR gains largely paralleled the effects seen at the SLR epoch, with an equivalently prominent impact of target direction. As mentioned above, all data in this section refers to stretch of the homonymous muscle i.e., all data originates from trials where the hand was perturbed along the direction of targets associated with stretch of the homonymous muscle, regardless of voluntary movement intent. Moreover, for the purposes of the LLR analyses, all load conditions were examined. That is, because goal-directed modulation of LLR gains is known to be robust against automatic gain-scaling, here we used the full ANOVA design of 2(preparatory delay) x 3(load) x 2(target distance) x 2(target direction). As expected (Fig. 8), LLR responses of all three muscles were strongly modulated as a function of target direction, with higher stretch reflex gains evident when preparing to reach targets associated with shortening of the homonymous muscle.

**Fig. 8.**
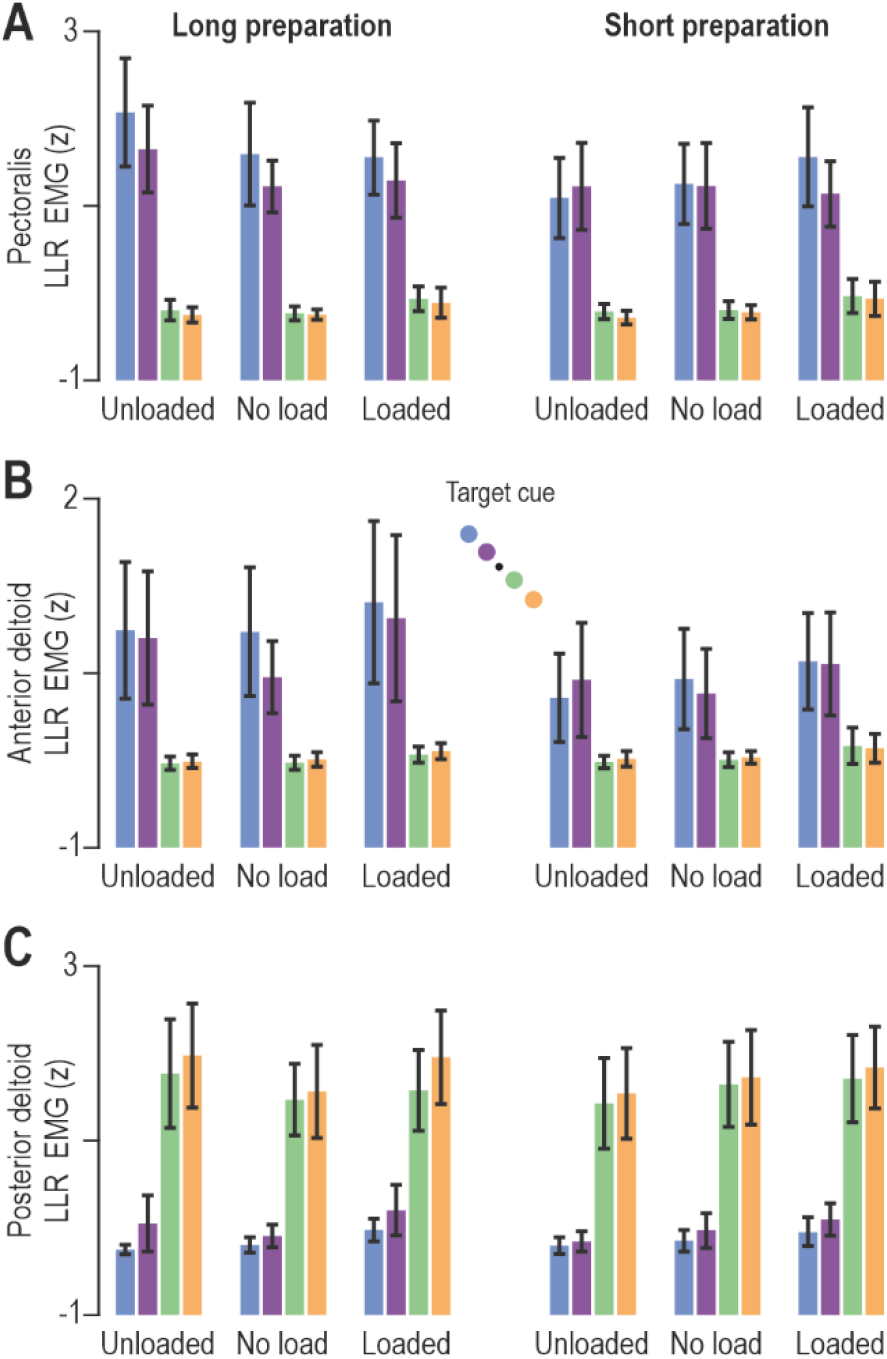
Goal-directed modulation of long-latency reflex gains. **A)** The coloured bars represent mean pectoralis LLR EMG (z) across participants (N=14), and vertical lines represent 95% confidence intervals. Colour-coding as in previous figures (see also schematic in ‘B’). ANOVA analyses indicated a significant impact of target direction on pectoralis LLR gains but, in contrast with the SLR results (Fig. 6A), the impact of target direction was significant on LLR gains also when the preparatory delay was short. However, there was also an effect of preparatory delay on LLR gains. **(B)** As ‘A’ but representing the anterior deltoid muscle. Similar LLR modulation patterns were observed as for the pectoralis. **(C)** As ‘A’ but representing the posterior deltoid.

Specifically, for the pectoralis muscle (Fig. 8A), the ANOVA indicated a significant main effect of target direction on LLR responses (F_1,13_=77.2, p<10^-5^ and 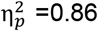), a significant main effect of preparatory delay (F_1,13_=5.8, p=0.032 and 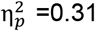), and a significant interaction between preparatory delay and target direction (F_1,13_=8.4, p=0.012 and 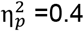). Tukey’s HSD indicated that all comparisons involving target direction and delay were significantly different (all p<0.009), with the exception of no significant effect of delay duration when preparing to reach a target in the direction of pectoralis lengthening (p=0.99; green and yellow bars, Fig 8A). In other words, delay duration played no role when the imposed pectoralis stretch was congruent with the goal of the intended movement. Instead, experiencing a long delay was associated with even stronger LLR responses when the hand was subsequently perturbed in the direction opposite to that of the intended reach (Fig 8A, blue and purple bars). Furthermore, there is a significant interaction effect between preparatory delay and load (F_2,26_=11.4, p=0.0003 and 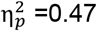) and a further interaction effect between delay, load and target direction (F_2,26_=6.1, p=0.007 and 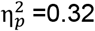; there is no main effect of load: F_2,26_=1.8, p=0.36). Post-hoc analyses revealed that LLR EMG was highest when the pectoralis was unloaded and the hand was perturbed in an incongruent direction following a long preparatory delay (i.e., left-most blue and purple bars in Fig. 8A; all p<0.0017). Hence, paralleling the SLR tuning of this muscle, goal-directed tuning of LLR was most prevalent when the unloaded pectoralis was perturbed/stretched following a long-enough preparatory delay (>250 ms). There is a significant but weak effect of target direction on LLR responses (F_1,13_=5, p=0.043 and 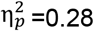) and a strong interaction effect between target distance and target direction (F_1,13_=10.2, p=0.007 and 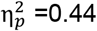). Tukey’s HSD indicated significantly higher LLR EMG for ‘far’ vs ‘near’ targets only when these were placed along the pectoralis shortening direction (i.e., blue vs. purple: p=0.01; green vs. orange targets: p=0.91; Fig. 8A).

Similar results were obtained for the anterior deltoid (Fig. 8B). An ANOVA indicated a significant main effect of target direction on LLR responses (F_1,13_=19.2, p=0.0008 and 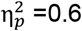), a significant main effect of preparatory delay (F_1,13_=12.3, p=0.004 and 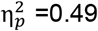) and an interaction effect between preparatory delay and target direction (F_1,13_=13.9, p=0.0026 and 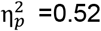). Tukey’s HSD test showed that all comparisons involving target direction and delay were significantly different (all p<0.0016), with the exception of no significant effect of delay duration when preparing to reach a target in the direction of pectoralis lengthening (p=0.98; green and yellow bars, Fig 8B). Like the case of the pectoralis, preparing to voluntarily shorten (vs. lengthen) the homonymous muscle is associated with stronger anterior deltoid LLRs, and this enhancement of LLR gains is even stronger following a long preparatory delay (Fig 8B, blue and purple bars in left panel vs right panel). Interestingly, like the case for SLR responses of this muscle, there is no significant impact of target distance on the LLR responses of the anterior deltoid (p=0.064). However, there is an interaction effect between preparatory delay and target distance (F_1,13_=7.4, p=0.018 and 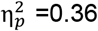), with post-hoc analysis showing that all relevant comparisons were significantly different (all p<0.003), with the exception of no significant impact of target distance when the delay was short (p=0.099). There is also a weak but significant main effect of load (F_2,26_=3.7, p<0.04 and 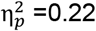), an interaction effect between load and target distance (F_2,26_=4.7, p<0.018 and 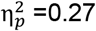), and a further interaction effect between load, target distance and direction distance (F_2,26_=4.9, p=0.017 and 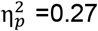). Tukey’s HSD test indicated a significant difference (p=0.00014) as a function of target distance only when preparing muscle shortening in the ‘no load’ condition (blue versus purple targets in the ‘no-load’ condition, Fig. 8B).

Equivalent results were obtained with regard to the posterior deltoid (Fig. 8B). The ANOVA indicated a significant main effect of target direction on posterior deltoid LLR (F_1,13_=78, p<10^-5^ and 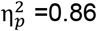) again demonstrating that the preparation to move in one direction or another sets up different feedback gains. There is an interaction effect between target distance and direction (F_1,13_=8.7, p=0.012 and 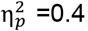). However, Tukey’s HSD only indicated a significant impact of target direction i.e., relative upregulation of LLR gains regardless whether the ‘shortening’ target was ‘near’ or ‘far’ (both p<0.0003). There is a weaker but significant main effect of load on posterior deltoid LLR responses (F_2,26_=4.4, p=0.023 and 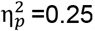) and an interaction effect between load and preparatory delay (F_2,26_=5.9, p=0.008 and 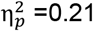). Tukey’s HSD test indicated that, a significant difference in LLR as a function of delay (i.e., higher LLR with longer delay) materialized only when the posterior deltoid was unloaded (p=0.034). There is also an interaction effect between load, preparatory delay and target direction (F_2,26_=3.9, p=0.032 and 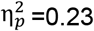), with post-hoc analyses indicating that the aforementioned impact of preparatory delay manifested only when preparing to shorten the unloaded posterior deltoid (p=0.0036). Overall, therefore, there is a consistent pattern of LLR goal-directed tuning: relatively weaker/stronger reflex gains when preparing to reach targets requiring stretch/shortening of the homonymous muscle, with an amplification of this effect in the unloaded muscle if a sufficiently long preparatory delay is allowed.

## Discussion

The aim of our study was to investigate the goal-directed tuning of stretch reflex gains in the context of delayed reaching, and further examine how such tuning depends on external loads and preparation duration. Participants were presented with 4 different targets (2 directions and 2 distances) under three different background loads. At two delays (short and long) after the target was presented to the participants, a rapid position-dependent perturbation was applied to the arm in order to test whether stretch reflex gains were changed in preparation for the upcoming reaching movement. Despite identical background loads and perturbations to the dominant upper limb, both the short and long latency stretch reflex responses (‘SLR’ and ‘LLR’) showed strong goal-directed modulation. Specifically, while target distance only produced small variations in the LLR gains, the direction of the prepared movement produced profound scaling of these responses. Moreover, as the delay between the target presentation and the perturbation increased, the LLR responses showed stronger changes according to the target direction, especially when the muscles were unloaded (i.e., when an external force was first applied in the direction of homonymous muscle action). This long delay condition also revealed a consistent strong goal-directed tuning of SLR gains, particularly in the unloaded condition. As with the LLR, we demonstrate that the SLR shows a clear modulation according to the presented target direction in all three analysed muscles, increasing in gain when the muscles will be shortened by the upcoming movement, and decreasing when they are expected to be lengthened. Overall, we show that preparing to reach a target produces a congruent modulation of the stretch reflex responses at both the SLR and LLR epochs, suggesting the independent involvement of the fusimotor system in movement preparation. A long-enough preparation (>250 ms) and muscle unloading (assistive loading) appear to trigger or enhance this independent preparatory control of reflex muscle stiffness (Figs. 6–7).

### SLR responses

With the exception of long-term reward-based training (Wolpaw, 1982), most previous research examining the modulation of SLR responses has shown little task-relevant modulation of feedback gains. In particular, until recently, SLR responses were considered to mainly exhibit gain-scaling or presynaptic modulation according to tasks. Gain scaling is the modulation of the feedback responses with the background activity or load (Matthews, 1986; Pruszynski et al., 2009). In addition to gain scaling, several papers have provided evidence using H-reflexes that pre-synaptic inhibition on the Ia afferents can affect the SLR response (Stein 1995; Capaday & Stein 1987; Nielsen & Kagamihara 1993; Perez et al., 2005). In general, these studies have shown modulation of the H-reflex for different tasks such as standing or walking, or during co-contraction, independent of the specific level of muscle activity. The modulation of the H-reflex during such tasks suggested a descending modulation of the SLR from pre- synaptic inhibition. More recently, Weiler and colleagues (Weiler et al., 2019; Weiler et al., 2021) have demonstrated that wrist posture influences the SLR responses at the elbow in a task-relevant manner, suggesting that spinal circuits are capable of integrating information from nearby joints. In a similar manner, although gain scaling is normally described within a single muscle or motor neuron pool, a recent study has shown SLR modulation that depends also on synergistic muscle activity across the shoulder joint (Nicolozakes et al., 2022). This modulation of SLR in the absence of movement or movement planning is compatible with the antagonistic muscle balance hypothesis (Dimitriou 2014), which has been supported independently (Villamar et al., 2022). Finally, it has also been recently shown that, despite the same initial posture, there is a goal-directed modulation of SLR gains when preparing to reach with the dominant limb (Papaioannou & Dimitriou, 2021). Specifically, when the perturbation stretching the homonymous muscle occurs in the direction of the cued target, there is a smaller SLR than if the same perturbation acts in the direction opposite to the target. Here we confirm this initial finding, showing again that the SLR is consistently modulated according to target direction.

In the absence of equivalent differences in pre-perturbation muscle activity, the effect of target direction on SLR suggests that movement preparation affects muscle spindle tuning, changing the SLR feedback gain during preparation via the independent control of fusimotor neurons. Indeed, SLR modulation across muscles relied on there being sufficient time between the target presentation and the perturbation, longer than the minimum time required for shaping reflex responses via selective CNS processing of sensory signals (e.g., Scott, 2016). In turn, this demonstrates that the preparatory modulation of reflex muscle stiffness requires sufficient time to completely unfold (>250 ms), which fits with the known slow-evolving nature of net fusimotor impact on spindle afferent responses (e.g., Crowe & Matthews, 1964a; Crowe & Matthews, 1964b). The proposal that independent fusimotor control is involved in movement preparation is also compatible with the demonstrated preparatory changes in the somatosensory cortex (Ariani et al., 2022) and the notion that feedback controllers are ‘loaded’ prior to movement onset (Ahmadi-Pajouh et al., 2012).

While we suggest that the goal-directed modulation of the SLR in our movement preparation task occurs through independent fusimotor control, there is still a possibility that the observed changes occurred through pre-synaptic inhibition. While this cannot be ruled out with the current study, we believe that this is unlikely for several reasons. First, presynaptic inhibition would be expected to provide consistent modulation of the SLR regardless of the background activity of the muscle, whereas in our study this did not occur when the muscle was directly loaded. Second, there is already evidence for a goal-directed preparatory modulation of muscle spindles in delayed reach (Papaioannou and Dimitriou, 2021), and more importantly this modulation matches the temporal evolution of the observed SLR tuning, where stronger modulation is found for delays larger than 250 ms. We therefore propose that the goal-directed tuning of SLR gain arises primarily through changes in gamma drive. The classic view that muscle spindle sensitivity relates to α-γ-coactivation (Vallbo, 1970; Vallbo et al., 1979) is changing to a more dynamic view where the fusimotor system allows flexible signal-processing at the periphery (Dimitriou, 2022). We find stronger goal-directed modulation of SLR responses when the muscle is unloaded. We hypothesize this arises because antagonist loading is accompanied by top-down reciprocal inhibition of lower motor neurons of the muscle, including gamma motor neurons (Dimitriou, 2014). The stronger goal-directed effects then arise as the independent goal-directed control of dynamic gamma motor neurons occurs on top of this blanket reciprocal inhibition of lower motor neurons that accompanies muscle unloading.

### LLR responses

It is well-known that LLR responses can vary according to task goals (e.g., Akazawa et al., 1983; Kimura et al., 2006; Nashed et al., 2012; Pruszynski et al., 2008). Early work demonstrated that simple commands to the participants such as to ‘relax’ or ‘resist’ the perturbation produced large variations in reflex responses (Crago et al., 1976; Rothwell et al., 1980). This was further refined by placing different targets either in the direction of the perturbation or the opposite direction in order to see whether the stretch reflex responses would be modified by the location and distance of the target (Pruszynski et al., 2008). The long latency responses were shown to modulate strongly according to the target location, increasing when the perturbation was in the opposite direction to the target and decreasing when perturbed in the direction of the target. Accordingly, it was recently shown that LLR gains are modulated when preparing to reach towards a cued target (Papaioannou & Dimitriou, 2021). In the present study, it was examined how these responses are affected by movement distance. Here, we found that the LLR responses showed strong modulation according to the direction of the planned future movement. That is, if the perturbation was in the direction opposite to the target of the upcoming movement, the LLR gain was increased, but decreased if the perturbation was in the direction of the target (Fig 7). When the delay between the target presentation and the perturbation was relatively short (250 ms), there was little effect of target distance or background load on the responses. However, additional preparation time (i.e., the long delay) was associated with stronger goal-directed modulation of LLR responses according to both target distance (especially for perturbations opposite to the target direction) and background load.

### Effect of preparatory delay

In our study there are strong differences according to the preparatory delay between target presentation and perturbation, both in the SLR and LLR epochs. Goal-directed differences at the SLR epoch were consistently observed only following the relatively long preparatory delay (Fig. 6). In the LLR epoch, for short preparatory delays we find clear tuning of the reflex gains according to the target direction, but there is little to no effect of target distance or background load. In contrast, for longer delays we find tuning of the feedback gains according to the target distance and modulation according to the background load, and an even stronger impact of target direction (Fig. 7). This goal directed modulation of feedback gains is appropriately tuned for the differences in the goals. That is, targets that are further away require stronger feedback gains. The difference in response for short and long delay conditions suggests that sufficient time is needed to determine how the feedback gains should modulate for further targets and different environmental dynamics. However, the current study only examined the stretch reflex modulation for two different delays (250 ms and 750 ms); we cannot define the minimum time required for full expression of goal-directed tuning of stretch reflexes. Further studies are needed to determine in more detail how such reflex modulation evolves over preparatory time.

Overall, we find larger differences in the goal directed feedback gains for longer preparatory delays. However, even more interesting is the difference between the LLR and SLR tuning as a function of preparatory delay. Our results show that even the relatively short delay of 250 ms was sufficient to systematically evoke tuning of the LLR responses according to target direction. However, no consistent modulation of the SLR was found for target direction or distance at this short delay. Instead, only for the longer delay did we find evidence of SLR modulation as a function of target direction. The above, and the presence of a stronger effect of target direction on LLR gains following a long delay, support the proposal that a slower-evolving mechanism (i.e. the independent fusimotor control of spindles) is also involved in the preparatory modulation of stretch reflex gains. Indeed, it has already been shown that muscle spindle Ia signals (and hence their fusimotor control) can affect LLR responses (e.g., Fellows et al., 1993; Hunter et al., 1988) that are likely mediated by both spinal and supraspinal circuits (Cheney & Fetz, 1984; Kurtzer, 2014; Pruszynski et al., 2011a; Soteropoulos & Baker, 2020).

### Effect of background loading

Many previous studies of stretch reflex modulation pre-loaded the muscles before applying perturbations. That is, they provide a background load to excite the muscle that they will stretch in order to elicit strong EMG responses. However, it can be seen in our study that such pre-loading to increase muscle activity also strongly affects the degree of reflex modulation. Specifically, there was no goal-directed modulation of SLR observed when the muscle was heavily loaded. This is likely due to the automatic gain-scaling feature which tends to saturate the SLR response (Matthews, 1986; Pruszynski et al., 2009). However, in the no-load or unloaded conditions we find goal-directed modulation of the SLR responses. These conditions reflect the everyday case of reaching to grasp an object, where the muscles are unloaded prior to the movement initiation. In contrast, pre-loading a muscle increases the alpha motor neuron drive, exciting the motoneuron pools, and appears to limit our ability to see such goal directed modulation at the SLR epoch. We suggest that it is critical for future studies to be performed under a range of loaded and unloaded conditions in order to examine the true range of feedback modulation. Moreover, the effects of background loading are even apparent within the LLR interval. In particular, we found the strongest goal-directed modulation of LLR and SLR responses to occur when the muscle was unloaded i.e., in cases where a background -‘assistive’- load was applied in the direction of homonymous muscle action. That goal-directed tuning of reflex muscle stiffness is enhanced by muscle unloading may partly account for the favourable impact of assistive loading for the purposes of motor rehabilitation (e.g., Wu et al., 2014).

### Effect of target parameters

In the current study we examine the goal-oriented modulation of stretch reflex responses by examining the effect of target distance. There is a long-standing history of examining changes in muscle activation with target distance (Buneo et al.,1994; Kaminski et al.,1995; Tyler & Karst, 2004), with timing and intensity varying as the distance increases. Scaling is also found for visuomotor responses, with variations in feedback gains to visual perturbations of the cursor with distance (Dimitriou et al., 2013) that can be explained by the time required to reach the target (Česonis & Franklin, 2020; Česonis & Franklin, 2022). However, there is very limited information regarding how target distance influences stretch reflex responses. In one study (Pruszynski et al., 2008), target distance was used to control the degree of resistance to the perturbations, showing a strong increase in LLR stretch gains with target distance for elbow movements.

Here we show similar goal-directed modulation of the LLR reflexes, particularly when the muscles are stretched against the direction of the target. However, we find that there is little or no variation with target distance when the homonymous muscle is lengthened in the direction of the cued target. Interestingly, we also find evidence for some target distance modulation of the SLR reflex when the muscles are stretched after longer preparatory periods. Nevertheless, the demonstrated impact of target direction on SLR gains strongly suggests that goal-directed tuning of spindles is a basic component of reach preparation (Papaioannou & Dimitriou, 2021). That is, reach preparation includes implementation of a plan at the periphery (i.e. modification of spindle gains), and does not only involve motor planning or otherwise priming of the CNS. In terms of the effect of target direction on reflex gains (relative down-regulation if the reach involves stretch of the homonymous muscle), it was more universal in our experimental results and shown in all three muscles. This result was rather easy to interpret as facilitating goal-directed behaviour, showing the compliance along the desired movement direction. However, at the same time we also found quite a robust effect of target distance on the pectoralis major SLR, which showed higher gains for ‘far’ targets, but this effect was not consistent across muscles.

### Reflex modulation and stiffness

The stiffness or compliance of the body has long been acknowledged as an important factor in motor control (Franklin & Wolpert, 2011; Hogan, 1984; McIntyre et al., 1995; Wolpert & Flanagan, 2010), but most reaching studies examine changes due to co-contraction (Franklin et al., 2007; Franklin & Franklin, 2021; Osu et al., 2002) or changing posture (Franklin et al., 2013; Lametti & Ostry, 2010; Trumbower et al., 2009). However, reflexes also contribute significantly to the stiffness of the muscles and limbs (Akazawa et al., 1983; Crago et al., 1976; Hoffer & Andreasse,n 1981; Kearney et al., 1997; Nichols & Houk, 1976). While stiffness due to co-contraction or posture provides an instantaneous response to a perturbation, reflexive contributions to stiffness act at a delay due to neural transmission delays and electromechanical delays of muscles (e.g., Ito et al., 2004). Due to this additional delay, reflex modulation is often thought to contribute more to the stiffness and stability in body posture tasks (Loram et al., 2011) than in object manipulation (Morasso, 2011), although both contribute (Franklin et al., 2007). Here we show evidence that there can be independent control of stretch reflexes via the fusimotor system, meaning that there can be task-dependent control of the rapid SLR for delayed reaching. These responses can modulate the force responses to perturbations quickly, as they do not have to wait for sensory input to reach the brain, be selectively processed in the brain, and then be passed back to the muscles. Indeed, independent fusimotor control provides true online goal-directed stiffness control, and with much lower energy consumption than co-contraction. As exemplified by the effect of preparatory delay in our study, such modulation does require some time to prepare, due at least in part to the slower nature of the gamma fusimotor system.

### Summary

In the current study we sought to examine the preparatory modulation of short- and long-latency stretch reflex responses in the dominant upper limb. While target distance was associated with relatively small variations in reflex gains, both short- and long-latency gains were strongly modulated as a function of target direction, in a manner that facilitated the upcoming voluntary movement. This goal-directed tuning of reflex gains was triggered or enhanced when the preparatory delay was sufficiently long (>250 ms) and the homonymous muscle was unloaded i.e., when a background load was first applied in the direction of homonymous muscle action (assistive loading). The results support the proposal that reach preparation also involves the goal-directed modulation of reflexive stiffness, likely via the independent control of fusimotor neurons.

## References

Ahmadi-Pajouh MA, Towhidkhah F, Shadmehr R (2012) Preparing to reach: selecting an adaptive long-latency feedback controller. J Neurosci 32(28): 9537–45. doi: 10.1523/JNEUROSCI.4275-11.2012

Akazawa K, Milner TE, Stein RB (1983) Modulation of reflex EMG and stiffness in response to stretch of human finger muscle. J Neurophysiol 49(1): 16–27. doi: 10.1152/jn.1983.49.1.16

Ariani G, Pruszynski JA, Diedrichsen J (2022) Motor planning brings human primary somatosensory cortex into action-specific preparatory states. Elife 11:e69517. doi: 10.7554/eLife.69517

Batista AP, Santhanam G, Yu BM, Ryu SI, Afshar A, Shenoy KV (2007) Reference Frames for Reach Planning in Macaque Dorsal Premotor Cortex. J Neurophysiol 98(2): 966–83. doi: 10.1152/jn.00421.2006

Buneo CA, Soechting JF, Flanders M (1994) Muscle activation patterns for reaching: the representation of distance and time. J Neurophysiol 71(4): 1546–58. doi: 10.1152/jn.1994.71.4.1546

Brooks, VB (1979) Motor programs revisited. In: Posture and Movement. (Talbot, RE; Humphrey, DR, eds.), pp. 13–49 New York: Raven Press

Capaday C, Stein RB (1987) Difference in the amplitude of the human soleus H reflex during walking and running. J Physiol 392:513–22. doi: 10.1113/jphysiol.1987.sp016794.

Česonis J, Franklin DW (2020) Time-to-Target Simplifies Optimal Control of Visuomotor Feedback Responses. eNeuro 7(2):ENEURO.0514-19.2020. doi: 10.1523/ENEURO.0514-19.2020

Česonis J, Franklin DW (2022) Contextual cues are not unique for motor learning: Task-dependant switching of feedback controllers. PLoS Comput Biol 18(6): e1010192. doi: 10.1371/journal.pcbi.1010192

Cheney PD, Fetz EE (1984) Corticomotoneuronal cells contribute to long-latency stretch reflexes in the rhesus monkey. J Physiol 349: 249–72. doi: 10.1113/jphysiol.1984.sp015155

Churchland MM, Cunningham JP, Kaufman MT, Foster JD, Nuyujukian P, Ryu SI, Shenoy KV (2012) Neural population dynamics during reaching. Nature 487: 51–6. doi: 10.1038/nature11129

Churchland MM, Cunningham JP, Kaufman MT, Ryu SI, Shenoy KV (2010) Cortical preparatory activity: representation of movement or first cog in a dynamical machine? Neuron 68(3): 387–400. doi: 10.1016/j.neuron.2010.09.015

Churchland MM, Santhanam G, Shenoy KV (2006) Preparatory activity in premotor and motor cortex reflects the speed of the upcoming reach. J Neurophysiol 96(6): 3130–46. doi: 10.1152/jn.00307.2006

Cluff T, Scott SH (2015) Apparent and Actual Trajectory Control Depend on the Behavioral Context in Upper Limb Motor Tasks. J Neurosci 35(36): 12465–76. doi: 10.1523/JNEUROSCI.0902-15.2015

Corneil BD, Olivier E, Munoz DP (2004) Visual responses on neck muscles reveal selective gating that prevents express saccades. Neuron 42(5):831–41. doi: 10.1016/s0896-6273(04)00267-3

Crago PE, Houk JC, Hasan Z (1976) Regulatory actions of human stretch reflex. J Neurophysiol 39(5): 925–35. doi: 10.1152/jn.1976.39.5.925

Crowe A, Matthews PB (1964a) The Effects of Stimulation of Static and Dynamic Fusimotor Fibres on the Response to Stretching of the Primary Endings of Muscle Spindles. J Physiol 174(1): 109–31. doi: 10.1113/jphysiol.1964.sp007476

Crowe A, Matthews PB (1964b) Further Studies of Static and Dynamic Fusimotor Fibres. J Physiol 174(1): 132–51. doi: 10.1113/jphysiol.1964.sp007477

Dimitriou M (2014) Human muscle spindle sensitivity reflects the balance of activity between antagonistic muscles. J Neurosci 34(41): 13644–55. doi: 10.1523/JNEUROSCI.2611-14.2014

Dimitriou M (2016) Enhanced Muscle Afferent Signals during Motor Learning in Humans. Curr Biol 26(8): 1062–8. doi: 10.1016/j.cub.2016.02.030

Dimitriou M (2018) Task-dependent modulation of spinal and transcortical stretch reflexes linked to motor learning rate. Behav Neurosci 132(3): 194–209. doi: 10.1037/bne0000241

Dimitriou M (2021) Crosstalk proposal: There is much to gain from the independent control of human muscle spindles. J Physiol 599(10): 2501–04. doi: 10.1113/JP281338

Dimitriou M (2022) Human muscle spindles are wired to function as controllable signal-processing devices. eLife 11: e78091. doi: 10.7554/eLife.78091

Dimitriou M, Franklin DW, Wolpert DM (2012) Task-dependent coordination of rapid bimanual motor responses. J Neurophysiol 107(3): 890–901. doi: 10.1152/jn.00787.2011

Dimitriou M, Wolpert DM, Franklin DW (2013) The temporal evolution of feedback gains rapidly update to task demands. J Neurosci 33(26): 10898–909. doi: 10.1523/JNEUROSCI.5669-12.2013

Fellows SJ, Domges F, Topper R, Thilmann AF, Noth J (1993) Changes in the short- and long-latency stretch reflex components of the triceps surae muscle during ischaemia in man. J Physiol 472: 737–48. doi: 10.1113/jphysiol.1993.sp019970

Franklin DW, Liaw G, Milner TE, Osu R, Burdet E, Kawato M (2007) Endpoint stiffness of the arm is directionally tuned to instability in the environment. J Neurosci 27(29): 7705–16. doi: 10.1523/JNEUROSCI.0968-07.2007

Franklin DW, Selen LP, Franklin S, Wolpert DM (2013) Selection and control of limb posture for stability. Conf Proc IEEE Eng Med Biol Soc 2013 2013: 5626–9. doi: 10.1109/EMBC.2013.6610826

Franklin DW, Wolpert DM (2011) Computational mechanisms of sensorimotor control. Neuron 72(3): 425–42. doi: 10.1016/j.neuron.2011.10.006

Franklin S, Franklin DW (2021) Feedback Gains modulate with Motor Memory Uncertainty. Neurons, Behavior, Data analysis, and Theory. 5(2):1–28. https://doi:10.51628/001c.22336

Ghez C, Favilla M, Ghilardi MF, Gordon J, Bermejo R, Pullman S (1997) Discrete and continuous planning of hand movements and isometric force trajectories. Exp Brain Res 115(2): 217–33. doi: 10.1007/pl00005692

Green DM, Swets JA (1966) Signal Detection Theory and Psychophysics. New York: Wiley.

Hammond PH (1956) The influence of prior instruction to the subject on an apparently involuntary neuro-muscular response. J Physiol 132(1): 17–8p

Hocherman S, Wise SP (1991) Effects of hand movement path on motor cortical activity in awake, behaving rhesus monkeys. Exp Brain Res 83(2): 285–302. doi: 10.1007/BF00231153

Hoffer JA, Andreassen S (1981) Regulation of soleus muscle stiffness in premammillary cats: intrinsic and reflex components. J Neurophysiol 45(2): 267–85. doi: 10.1152/jn.1981.45.2.267

Hogan N (1984) An organizing principle for a class of voluntary movements. J Neurosci 4(11): 2745–54. doi: 10.1523/JNEUROSCI.04-11-02745.1984

Hunter JP, Ashby P, Lang AE (1988) Afferents contributing to the exaggerated long latency reflex response to electrical stimulation in Parkinson’s disease. J Neurol Neurosurg Psychiatry 51(11): 1405–10. doi: 10.1136/jnnp.51.11.1405

Ito T, Murano EZ, Gomi H (2004) Fast force-generation dynamics of human articulatory muscles. J Appl Physiol (1985) 96(6): 2318–24; discussion 2317. doi: 10.1152/japplphysiol.01048.2003

Kaminski TR, Bock C, Gentile AM (1995) The coordination between trunk and arm motion during pointing movements. Exp Brain Res 106(3): 457–66. doi: 10.1007/BF00231068

Kearney RE, Stein RB, Parameswaran L (1997) Identification of intrinsic and reflex contributions to human ankle stiffness dynamics. IEEE Trans Biomed Eng 44(6): 493–504. doi: 10.1109/10.581944

Kimura T, Haggard P, Gomi H (2006) Transcranial magnetic stimulation over sensorimotor cortex disrupts anticipatory reflex gain modulation for skilled action. J Neurosci 26(36): 9272–81. doi: 10.1523/JNEUROSCI.3886-05.2006

Kurtzer IL (2014) Long-latency reflexes account for limb biomechanics through several supraspinal pathways. Front Integr Neurosci 8: 99. doi: 10.3389/fnint.2014.00099

Kutas M, Donchin E (1974) Studies of squeezing: handedness, responding hand, response force, and asymmetry of readiness potential. Science 186(4163): 545–8. doi: 10.1126/science.186.4163.545

Lametti DR, Ostry DJ (2010) Postural constraints on movement variability. J Neurophysiol 104(2): 1061–7. doi: 10.1152/jn.00306.2010

Lee H, Perreault EJ (2019) Stabilizing stretch reflexes are modulated independently from the rapid release of perturbation-triggered motor plans. Sci Rep 9(1): 13926. doi: 10.1038/s41598-019-50460-1

Loram ID, Gollee H, Lakie M, Gawthrop PJ (2011) Human control of an inverted pendulum: is continuous control necessary? Is intermittent control effective? Is intermittent control physiological? J Physiol 589(Pt 2): 307–24. doi: 10.1113/jphysiol.2010.194712

Marsden CD, Merton PA, Morton HB (1972) Servo action in human voluntary movement. Nature 238(5360): 140–3. doi: 10.1038/238140a0

Matthews PB (1986) Observations on the automatic compensation of reflex gain on varying the pre-existing level of motor discharge in man. J Physiol 374: 73–90. doi: 10.1113/jphysiol.1986.sp016066

McIntyre J, Gurfinkel EV, Lipshits MI, Droulez J, Gurfinkel VS (1995) Measurements of human force control during a constrained arm motion using a force-actuated joystick. J Neurophysiol 73(3): 1201–22. doi: 10.1152/jn.1995.73.3.1201

Messier J, Kalaska JF (2000) Covariation of primate dorsal premotor cell activity with direction and amplitude during a memorized-delay reaching task. J Neurophysiol 84(1): 152–65. doi: 10.1152/jn.2000.84.1.152

Morasso P (2011) ‘Brute force’ vs. ‘gentle taps’ in the control of unstable loads. J Physiol 589(Pt 3): 459–60. doi: 10.1113/jphysiol.2010.203604

Nashed JY, Crevecoeur F, Scott SH (2012) Influence of the behavioral goal and environmental obstacles on rapid feedback responses. J Neurophysiol 108(4): 999–1009. doi: 10.1152/jn.01089.2011

Nashed JY, Crevecoeur F, Scott SH (2014) Rapid online selection between multiple motor plans. J Neurosci 34(5): 1769–80. doi: 10.1523/JNEUROSCI.3063-13.2014

Nichols TR, Houk JC (1976) Improvement in linearity and regulation of stiffness that results from actions of stretch reflex. J Neurophysiol 39(1): 119–42. doi: 10.1152/jn.1976.39.1.119

Nicolozakes CP, Sohn MH, Baillargeon EM, Lipps DB, Perreault EJ (2022) Stretch reflex gain scaling at the shoulder varies with synergistic muscle activity. J Neurophysiol. 128(5):1244–1257. doi: 10.1152/jn.00259.2022

Nielsen J, Kagamihara Y (1993) The regulation of presynaptic inhibition during co-contraction of antagonistic muscles in man. J Physiol 464:575–93. doi: 10.1113/jphysiol.1993.sp019652

Osu R, Franklin DW, Kato H, Gomi H, Domen K, et al. (2002) Short- and long-term changes in joint co-contraction associated with motor learning as revealed from surface EMG. J Neurophysiol 88(2): 991–1004. doi: 10.1152/jn.2002.88.2.991

Papaioannou S, Dimitriou M (2021) Goal-dependent tuning of muscle spindle receptors during movement preparation. Sci Adv 7(9): eabe0401. doi: 10.1126/sciadv.abe0401

Perez MA, Lungholt BK, Nielsen JB (2005) Presynaptic control of group Ia afferents in relation to acquisition of a visuo-motor skill in healthy humans. J Physiol 568(Pt 1):343–54. doi: 10.1113/jphysiol.2005.089904

Pruszynski JA, Kurtzer I, Lillicrap TP, Scott SH (2009) Temporal evolution of “automatic gain-scaling”. J Neurophysiol 102(2): 992–1003. doi: 10.1152/jn.00085.2009

Pruszynski JA, Kurtzer I, Nashed JY, Omrani M, Brouwer B, Scott SH (2011a) Primary motor cortex underlies multi-joint integration for fast feedback control. Nature 478: 387–90. doi: 10.1038/nature10436

Pruszynski JA, Kurtzer I, Scott SH (2008) Rapid motor responses are appropriately tuned to the metrics of a visuospatial task. J Neurophysiol 100(1): 224–38. doi: 10.1152/jn.90262.2008.

Pruszynski JA, Kurtzer I, Scott SH (2011b) The long-latency reflex is composed of at least two functionally independent processes. J Neurophysiol 106(1): 449–59. doi: 10.1152/jn.01052.2010

Rothwell JC, Traub MM, Marsden CD (1980) Influence of voluntary intent on the human long-latency stretch reflex. Nature 286(5772): 496–8. doi: 10.1038/286496a0

Scott SH (2016) A Functional Taxonomy of Bottom-Up Sensory Feedback Processing for Motor Actions. Trends Neurosci 39(8): 512–26. doi: 10.1016/j.tins.2016.06.001

Shadmehr R, Krakauer JW (2008) A computational neuroanatomy for motor control. Exp Brain Res 185(3): 359–81. doi: 10.1007/s00221-008-1280-5

Shemmell J, Krutky MA, Perreault EJ (2010) Stretch sensitive reflexes as an adaptive mechanism for maintaining limb stability. Clin Neurophysiol 121(10): 1680–89. doi: 10.1016/j.clinph.2010.02.166

Soteropoulos DS, Baker SN (2020) Long-latency Responses to a Mechanical Perturbation of the Index Finger Have a Spinal Component. J Neurosci 40(20): 3933–48. doi: 10.1523/JNEUROSCI.1901-19.2020

Stein R.B (1995) Presynaptic inhibition in humans. Prog Neurobiol, 47(6), 533–544. doi: 10.1016/0301-0082(95)00036-4

Sternberg S, Monsell S, Knoll RL, Wright CE (1978) The latency and duration of rapid movement sequences: Comparisons of speech and typewriting. In: Information Processing in Motor Control and Learning (Stelmach, GE, ed). pp. 117–52. New York: Academic Press

Tanji J, Evarts EV (1976) Anticipatory activity of motor cortex neurons in relation to direction of an intended movement. J Neurophysiol 39(5): 1062–8. doi: 10.1152/jn.1976.39.5.1062

Todorov E, Jordan MI (2002) Optimal feedback control as a theory of motor coordination. Nat neurosci 5(11): 1226–35. doi: 10.1038/nn963

Trumbower RD, Krutky MA, Yang BS, Perreault EJ (2009) Use of self-selected postures to regulate multi-joint stiffness during unconstrained tasks. PLoS One 4(5): e5411. doi: 10.1371/journal.pone.0005411

Tyler AE, Karst GM (2004) Timing of muscle activity during reaching while standing: systematic changes with target distance. Gait Posture 20(2): 126–33. doi: 10.1016/j.gaitpost.2003.07.001

Vallbo AB (1970) Discharge patterns in human muscle spindle afferents during isometric voluntary contractions. Acta Physiol Scand 80(4):552–66. doi: 10.1111/j.1748-1716.1970.tb04823.x

Vallbo AB, Hagbarth KE, Torebjörk HE, Wallin BG (1979) Somatosensory, proprioceptive, and sympathetic activity in human peripheral nerves. Physiol Rev 59(4):919–57. doi: 10.1152/physrev.1979.59.4.919

Villamar Z, Ludvig D, Perreault EJ (2022) Short latency stretch reflexes depend on the balance of activity in agonist and antagonist muscles during ballistic elbow movements. J Neurophysiol Epub ahead of print. doi: 10.1152/jn.00171.2022.

Wagner MJ, Smith MA (2008) Shared Internal Models for Feedforward and Feedback Control. J Neurosci 28(42): 10663–73. doi: 10.1523/JNEUROSCI.5479-07.2008

Weiler J, Gribble PL, Pruszynski JA (2019) Spinal stretch reflexes support efficient hand control. Nat Neurosci 22: 529–33. doi: 10.1038/s41593-019-0336-0

Weiler J, Gribble PL, Pruszynski JA (2021) Spinal stretch reflexes support efficient control of reaching. J Neurophysiol 125(4): 1339–47. doi: 10.1152/jn.00487.2020

Wise SP (1985) The primate premotor cortex: past, present, and preparatory. Annu Rev Neurosci 8: 1–19. doi: 10.1146/annurev.ne.08.030185.000245

Wolpaw JR (1982) Change in short-latency response to limb displacement in primates. Fed Proc 41(6): 2156–9.

Wolpert DM, Flanagan JR (2010) Motor learning. Curr Biol 20(11): R467–72. doi: 10.1016/j.cub.2010.04.035

Wu M, Landry JM, Kim J, Schmit BD, Yen SC, Macdonald J (2014) Robotic resistance/assistance training improves locomotor function in individuals poststroke: a randomized controlled study. Arch Phys Med Rehabil 95(5): 799–806. doi: 10.1016/j.apmr.2013.12.021

Yang L, Michaels JA, Pruszynski JA, Scott SH (2011) Rapid motor responses quickly integrate visuospatial task constraints. Exp Brain Res 211(2): 231–42. doi: 10.1007/s00221-011-2674-3

Yeo SH, Franklin DW, Wolpert DM (2016) When Optimal Feedback Control Is Not Enough: Feedforward Strategies Are Required for Optimal Control with Active Sensing. PLoS Comput Biol 12(12): e1005190. doi: 10.1371/journal.pcbi.1005190

